# miRNA activity inferred from single cell mRNA expression

**DOI:** 10.1101/2020.07.14.202051

**Authors:** Morten Muhlig Nielsen, Jakob Skou Pedersen

## Abstract

High throughput single-cell RNA sequencing (scRNAseq) can provide mRNA expression profiles for thousands of cells. However, miRNAs cannot currently be studied at the same scale. By exploiting that miRNAs bind well-defined sequence motifs and typically down-regulate target genes, we show that motif enrichment analysis can be used to derive miRNA activity estimates from scRNAseq data.

Motif enrichment analyses have traditionally been used to derive binding motifs for regulatory factors, such as miRNAs or transcription factors, that have an effect on gene expression. Here we reverse its use. By starting from the miRNA seed site, we derive a measure of activity for miRNAs in single cells. We first establish the approach on a comprehensive set of bulk TCGA cancer samples (n=9,679), with paired mRNA and miRNA expression profiles, where many miRNAs show a strong correlation with measured expression. By downsampling we show that the method can be used to estimate miRNA activity in sparse data comparable to scRNAseq experiments. We then analyze a human and a mouse scRNAseq data set, and show that for several miRNA candidates, including liver specific miR-122 and muscle specific miR-1 and miR-133a, we obtain activity measures supported by the literature. The methods are implemented and made available in the miReact software. Our results demonstrate that miRNA activities can be estimated at the single cell level. This allows insights into the dynamics of miRNA activity across a range of fields where scRNAseq is applied.

## Introduction

The introduction of single-cell RNA sequencing (scRNAseq) is having an immense impact on our understanding of spatial and temporal gene expression dynamics and is allowing studies of gene expression in rare cell types. As most scRNAseq techniques rely on primer extension from the polyadenylated tail of messenger RNAs (mRNAs), they primarily quantify protein-coding transcripts. Other classes of transcripts and in particular microRNAs (miRNAs), cannot be studied as readily or at the same scale. Though progress has been made on measuring miRNA expression in single cells, original fluorescence-based methods can only evaluate a few miRNAs per cell^1–3^, while recent sequencing-based methods require extensive cell handling and thus have only been applied to few cells^4–7^. Single cell miRNA sequencing is thus still not standard practice.

Here we explore the potential to evaluate miRNA activity levels based on mRNA expression profiles. We exploit that miRNAs typically destabilise their targets and that each miRNA can bind to the 3’UTRs of many mRNAs. The activity of a miRNA can thus be evaluated statistically by considering the relative expression levels of all target genes.

We^8^ and others^9–13^ have previously developed statistical methods that exploit this idea to study sequence motifs. These approaches have typically identified motifs that associate with expression levels based on case and control experiments, e.g. with overexpression of a molecule of interest, such as a miRNA^9–13^. We here reverse this setup and exploit that the binding motifs (target sites) of miRNAs are known. We can thereby evaluate miRNA activity scores based on expression profiles across data sets with many samples.

We first demonstrate that miRNA activity scores, inferred from bulk mRNA expression profiles, correlate with bulk miRNA expression levels across a set of nearly ten thousand pan-cancer samples from The Cancer Genome Atlas (TCGA)^14^. These observations further extend to a set of nearly eight thousand normal tissue samples from GTEx^15^. By downsampling, we reveal a potential for using this approach in sparse scRNAseq datasets. Focussing on miR-122, which is known to be highly tissue-specific^16^, we provide a proof of principle for the use of miRNA activity scores in both mouse and human scRNAseq datasets and confirm that miR-122 is primarily active in liver, and at the single cell level, primarily in hepatocytes. We further show examples of other miRNAs with tissue and cell type specific activity scores.

As single cell miRNA sequencing remains challenging to perform at scale, the miReact approach offers an opportunity to study miRNA biology in large-scale single cell experiments, at no additional cost over scRNAseq. The power and scope to infer single cell miRNA activities from scRNAseq experiments will further benefit from a continued increase in read depth and cellular throughput.

## Results

### Inference of miRNA activity from mRNA expression profiles

We first sought to establish whether miRNA acitivity could be inferred from mRNA expression profiles from bulk-tissue RNAseq. For this, we extracted 9,679 mRNA expression profiles from TCGA cancer samples^14^, most of which had matched miRNA expression (n=9,366; Figure 1a). The basic idea of the approach is to evaluate if genes with miRNA target sites (binding motifs) are downregulated compared to other genes. To make expression levels comparable between genes, the expression levels are traditionally transformed into fold-changes based on a control sample, as used by related motif-analysis methods^8–10,13^. As we lack control samples in our setting, we calculate fold-changes relative to the median gene expression level across all samples (Figure 1b).

**Figure 1:**
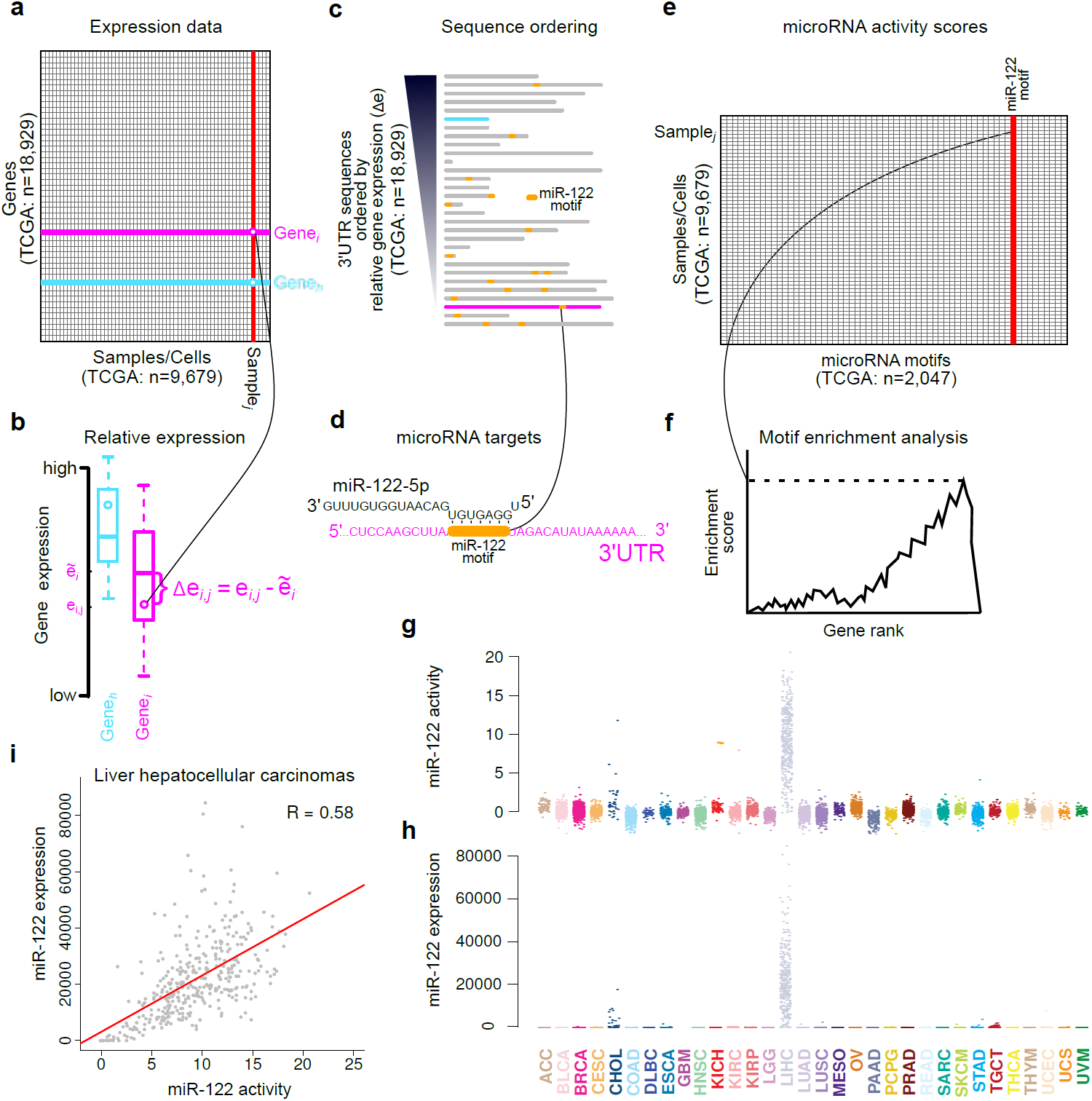
Method overview and illustration on miR-122 and TCGA data. Gene level expression data (**a**) is converted into sample/cell relative expression (Δe) (**b**) by subtraction of gene medians (denoted by ∼) obtained across all samples/ cells. For each sample/cell, 3’UTR sequences sorted by relative gene expression (**c**) are annotated withmiRNA motif presence (**d**) and subjected to motif enrichment analysis evaluating motifs’ association with expression (**f**). miRNA activity scores (**e**) are derived from the motif enrichment statistics to produce scores for each individual sample/cell for all miRNA binding motifs. **g**,**h**, Activity scores for miR-122-5p (**g**) and miR-122 expression (**h**) in TCGA samples. TCGA abbreviations are in Supplemental Table 1. **i**, Correlations between miR-122-5 activity scores and miR-122 expression across liver hepatocellular carcinomas (LIHC).

We thereafter evaluate the activity score for a comprehensive set of miRNAs^17^ (n=2,047) for each sample. For each miRNA we annotate the 3’UTR sequences with presence of binding sites. We exploit that miRNA binding is primarily defined by the 5’ seed site --a seven nucleotide long motif that has perfect complementarity to the miRNA targets^18^. miRNA binding models have proven efficient to evaluate activity of ectopically expressed miRNAs in cell line experiments in vitro^8–10,12,13^. For each sample, the 3’UTR sequences are ranked by their relative expression and the presence of motifs scored (Figure 1c). Finally, miRNA activity is statistically evaluated as the association of 3’UTR motif enrichments with gene expression, using either of three previously developed measures^8,9^ (Figure 1e,f). For further details, see Methods and Supplemental note.

### miRNA activity as a miRNA expression proxy

As a proof of concept, we first focussed on the activity scores for miR-122, which is an established liver specific miRNA. miR-122 has more activity in liver cancer samples than in samples with other tissues of origin (Figure 1g). This also corresponds well with the miR-122 expression levels observed across the same samples (Figure 1h), which indeed are positively correlated with the activity scores both in the liver cohort (R=0.58, p<2e-16, Figure 1i) and across all cancer types (R=0.85, p<2e-16, Figure 2a). This pattern is absent for the target of miR-122-3p, the opposite strand to the mature miRNA miR-122-5p, thus supporting that this approach can evaluate the activity of miRNAs in a strand specific manner (Supplemental Figure 1).

**Figure 2:**
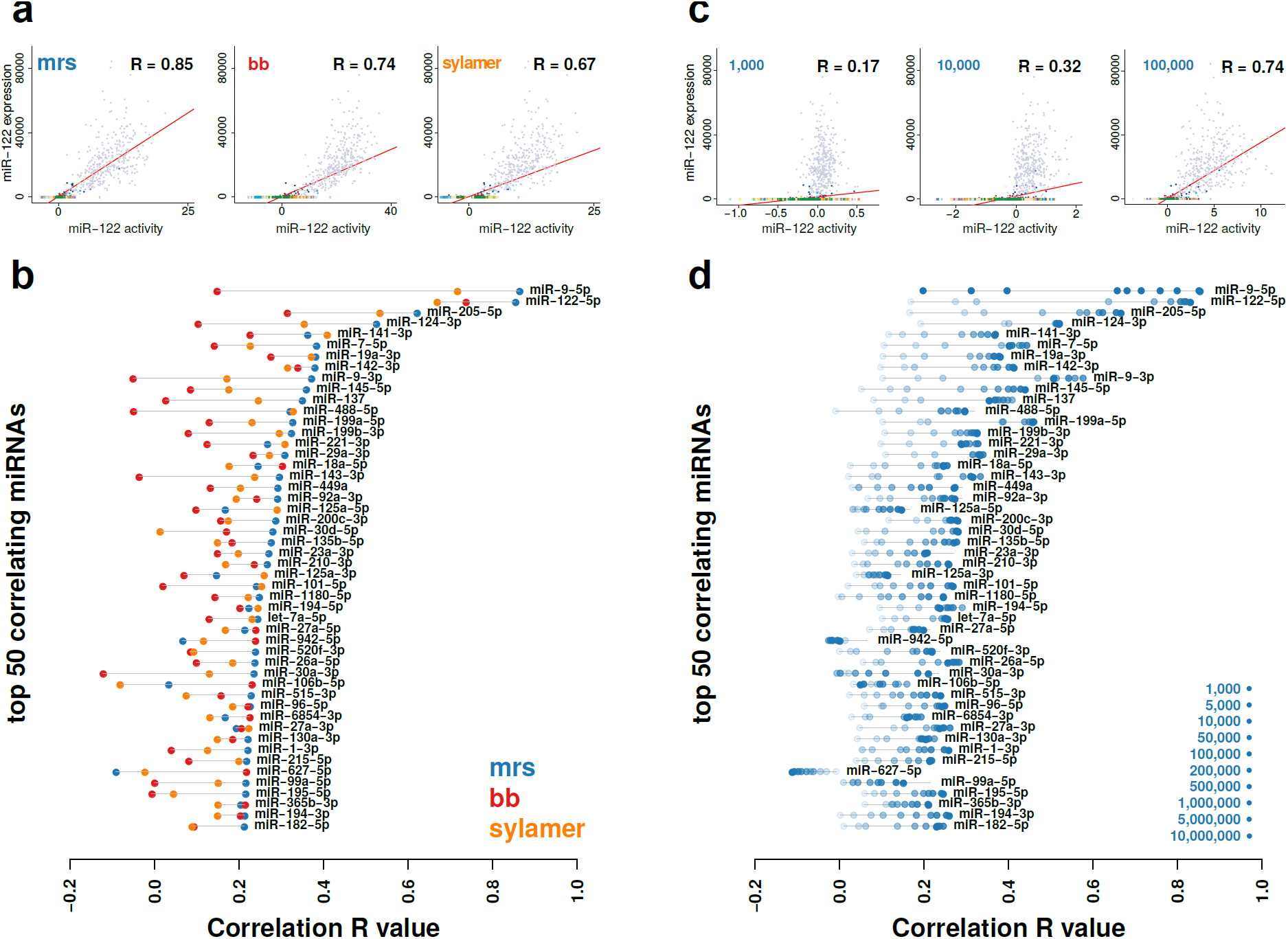
Method evaluation. **a**, (Top) Correlation of miR-122 activity with expression for the three different motif enrichment methods across the TCGA data set (n=9,679). (Bottom) Correlation values for the 50 miRNAs with highest correlation between activity and expression for any of the three activity score evaluation methods. **b**, (Top) plots as in (**a**) based on the mrs method for downsampled TCGA data sets with indicated number of total read counts per sample (shades of blue). (Bottom) Correlation values for top 50 miRNAs as in (**a**) based on the mrs method for 10 different downsampled TCGA data sets.

We next performed a more comprehensive evaluation of how different types of activity scores correlate with miRNA expression across the TCGA data set. More concretely, we compared three different ways of calculating motif enrichment scores. The first relies on a modified rank sum (mrs) statistic and the second on a brownian bridge (bb) statistic, both implemented in Regmex^8^. The third relies on the Sylamer method, which is based on a hyper geometric statistic^9^. For miR-122, all three approaches assign high scores to the liver cancer samples (Figure 1g and Supplemental Figure 2) and significant correlations between expression and activity, with the mrs performing best (Figure 2a). We then repeated this analysis across all miRNAs in our set and ranked them by their maximal correlation across all three ways of calculating activity scores. Among top-50, the mrs most often showed the highest correlation (Figure 2a, bottom). We thus used mrs for the remaining analysis. The full method is implemented and made available in the miReact tool.

### Cases of tissue specific miRNA activity

Having established that miRNA activities can be inferred from gene expression of TCGA samples, using miR-122 as a case study, we looked further into some of the other miRNAs with high activity scores and strong miRNA expression correlation (Figure 2b). We expanded the analysis to include the comprehensive set of healthy human tissue samples from the GTEx data set^15^, which again showed the expected liver-specific expression pattern for miR-122 (Supplemental Figure 3).

miR-9 and miR-124 are both top-ranked by miRNA expression correlation in TCGA (Figure 2b) and both known to be expressed in brain tissue^19,20^. They primarily show activity in gliomas (LGG) and glioblastomas (GBM, Supplemental Figure 4). The activity in the glioma samples are significantly correlated with miRNA expression. The glioblastoma samples do not have miRNA expression measures in the TCGA data set, however, based on these results, we hypothesize that miR-9 is expressed. Similar expression patterns are observed In the healthy brain tissue from GTEx (Supplemental Figure 3).

We then looked at miR-1, a muscle specific miRNA^21,22^, which shows activity in GTEx skeletal muscle and heart tissues, as we would also expect (Supplemental Figure 3). Finally, we looked at miR-7, a miRNA that was shown to bind to the circular RNA circ7as, which regulates its activity by acting as a miRNA sink^23,24^. Here we saw a higher activity in the pituitary gland, hypothalamus, and adrenal gland, in line with a neuroendocrine role for miR-7 in human, as has been established in fish, mouse, and pig^25–28^ (Supplemental Figure 3).

### Downsampling suggests robustness at the single cell level

The above results suggest that this approach might be used as an indicator of miRNA activity in single cell expression (scRNAseq) experiments. However, of primary concern is the data sparsity of such experiments. To investigate the potential, we created down-sampled versions of the TCGA data set. We assigned a total number of reads for each sample, and sampled reads proportional to each genes’ expression value, thus simulating a situation of sparsity. We then ran miReact for different numbers of total reads spanning from 1,000 to 10,000,000 reads per sample. scRNAseq experiments can provide meaningful cell type classification at library sizes above 10,000 reads^29,30^ and expression level saturation occurs at around 1,000,000 reads^31^.

We observed that even at 1,000 reads, we see positive, albeit small, correlations with expression for the top correlating miRNAs (Figure 2c,d, Supplemental Figure 5). Increasing the reads cause correlations to rapidly increase towards the values of the full TCGA expression set. In particular, going from 10,000 to 50,000 reads has the largest effect on the correlations for many miRNAs. There is thus reason to attempt to apply the method on scRNAseq expression data to see if already known findings can be reproduced.

### miR-122 activity in single cells

We first investigated miR-122 activity in two scRNAseq data sets: The ‘Tabula Muris’ of 20 mouse organs that contains 44,949 FACS sorted cells ^32^, which has a high read count (median > 400,000 reads/cell, see methods) and a human liver experiment with 10,372 cells ^33^ and a much lower read count (median < 3000 reads/cell). For the mouse data set, as with TCGA and GTEx, we saw that miR-122 activity was higher in liver tissue, and thus at the single cell level could reproduce this observation (Figure 3a-c).

**Figure 3:**
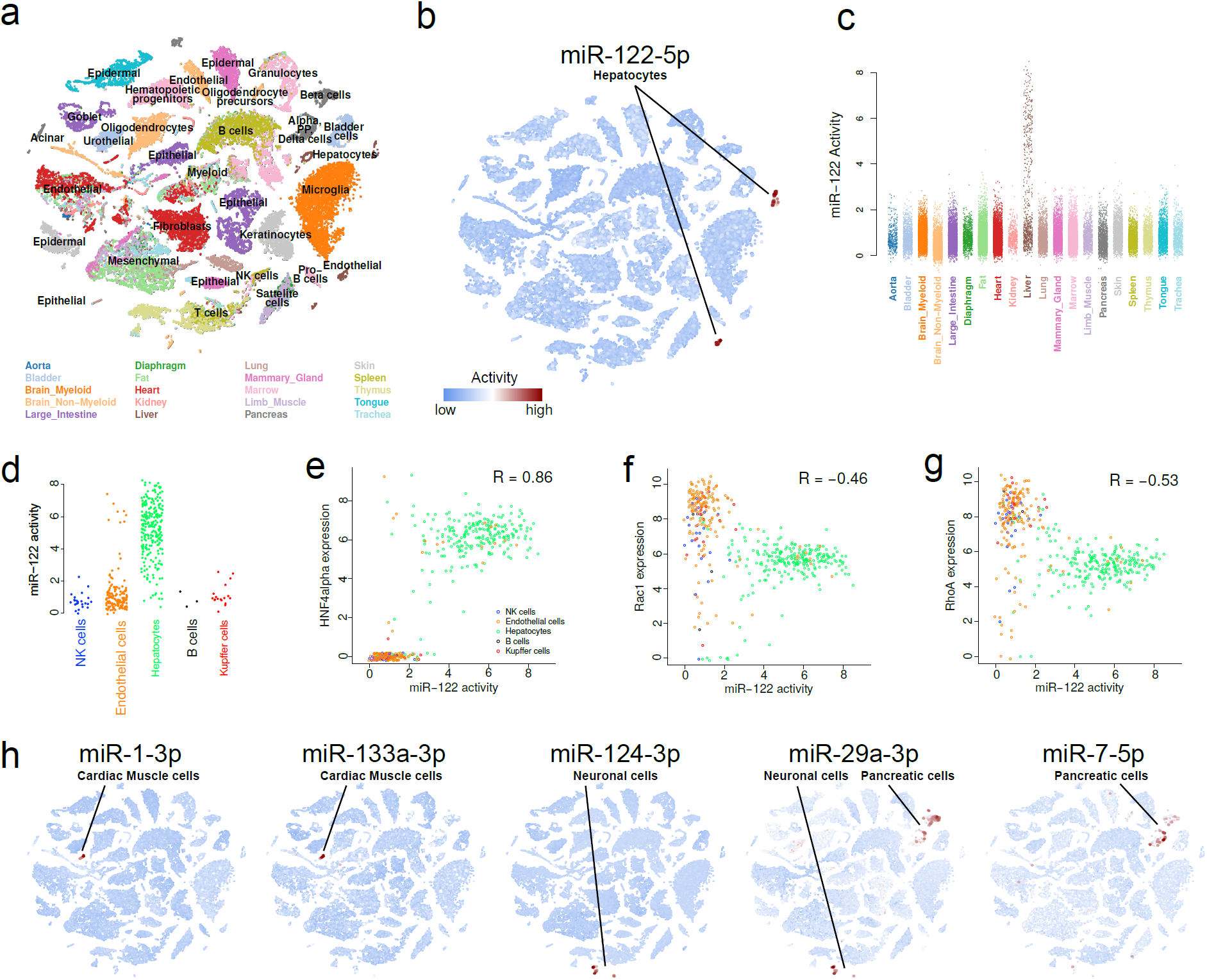
miRNA activity in single cells. **a**, t-SNE plot showing clusters of cell types colored by tissue types for 42,192 cells of the Tabula Muris data set. Embeddings and cell annotations were obtained from the original work. **b**, miR-122 activity overlaid the t-SNE embeddings as in (**a**) with indication of cell types for high activity cells. **c**, miR-122 activity of individual cells separated into tissues. **d**, miR-122 activity of individual liver cells separated into cell types. **e**,**f**,**g**, Correlation between mir-122 activity and expression of the miR-122 transcription factor HNF4alpha (**e**), and miR-122 targets Rac1(**f**), and RhoA (**g**). **h**, miRNA activities overlaid t-SNE embeddings as in (**a**) with indication of cell types for high activity cells. Cells with less than 2000 expressed genes were removed in (**d-g**).

Unlike the bulk liver samples, there were many mouse liver cells with low miR-122 activity. We further looked into this by sub-setting the liver cells based on the cell type annotations provided by the consortium^32^. Here we observed, as expected, that miR-122 activity mainly resides in hepatocytes, and correlates with expression of HNF4alpha, a transcription factor known to regulate miR-122^34,35^ (Figure 3d,e). Moreover, the expression of two known targets of miR-122, Rac1 and RhoA^35^, correlates negatively with the miRNA activity (Figure 3f,g). This thus supports that miRNA binding motif enrichment analysis may be useful for the study of miRNA activity at the single cell level. There were hepatocytes which appeared not to have any miR-122 activity. These cells had a low number of expressed genes (<2000), indicating a limit in sensitivity when expression data is sparse (Supplemental figure 6).

In the human liver data set, despite a 100 fold lower median read count, we also saw an increased miR-122 activity in hepatocytes as well as in EPCAM+ cells relative to other cell types, although here the signal was less pronounced (Supplemental Figure 7a-c). Again, there is a positive correlation with HNF4alpha expression as well as negative correlations with both target genes, but here as well, the effect was smaller than in the mouse data set (Supplemental Figure 7d-f). This is likely due to low read counts in some hepatocytes (Supplemental Figure 8).

### Cell type specific miRNA activity

Discovery of cell type specific miRNA activity in single cell data sets opens for analysis of miRNAs in rare cell types that can be difficult to obtain with traditional sampling techniques. We thus looked for additional examples of miRNAs with high activity in a single or few cell types in the mouse data set. By comparing the miRNA activity distributions for each cell type, scoring for cell types with outlier behaviour, we identified several miRNAs with cell type specific activity. Muscle specific miRNAs miR-1 and miR-133 were active in cardiac muscle cells, but had low activity in other cell types of the heart including smooth muscle cells (Figure 3, Supplemental Figure 9). miR-124 activity was specific to neuronal brain cells and miR-29a and miR-7 showed activity in pancreatic cells as well as neuronal cells. This generally fits with observations in the bulk data sets and the literature^19,36^, although we did not observe increased activity of miR-7 in pancreas samples from GTEx (Supplemental Figure 4). miR-7 has been reported to be active in pancreas in both mouse and human in a cell specific manner^37,38^. It is thus possible that a bulk experiment would conceal a signal from the target genes because of dilution by other cell types.

## Discussion

We have shown that motif enrichment analysis can be useful to estimate activity of miRNAs in high-throughput expression experiments, including scRNAseq. Measuring miRNA expression has often not been a priority in consortia, despite the importance of understanding these regulatory molecules. At the single cell level, current protocols lack the possibility of creating miRNA expression at a high-throughput scale. This method can thus be a useful tool for the study of miRNAs under conditions where expression measures are not easily obtainable.

Yet, far from all miRNAs show a good correlation between calculated activity and expression. A number of reasons may explain this. The activity measure is based on reduced expression of mRNAs with miRNA targets. However, besides leading to mRNA degradation, miRNAs are known to also exert their effect by inhibiting translation without reduction in mRNA expression^39^. Thus this effect may be absent from the activity signal. Other factors than seed site complementarity, e.g. local mRNA structure, can also be important for binding^18^. The 3’UTR model is based on the longest sequence among potentially multiple isoforms, some of which could be tissue specific, and thus the model for some genes in some samples or cells could be wrong. Finally, the model for the expected gene expression, the median across the expression set, does not accurately reflect gene expression in the absence of the miRNA, but rather is a surrogate. It has the advantage over merely using sample gene expression, that genes turned off in most samples will not be ranked extreme in general.

We note that for a miRNA to show good activity to expression correlation in a dataset, it needs to hold some variation in expression of a miRNA. This is not necessarily the case in the TCGA dataset. An example is miR-1, which is active in muscle tissue. Since there are no muscle-derived samples in the TCGA cancer set, we saw only a modest correlation between activity and expression. For the GTEx and mouse single cell data set, however, we saw a highly specific signal in muscle and heart tissue, clearly indicating that activity reflects expression of miR-1, although not demonstrable in the TCGA dataset.

Inferring miRNA activity may provide additional insight over measuring expression. Although we saw a good correlation between miR-7 activity and expression, miR-7 sequestration into a complex with the circular RNA circ7as^23,24^ may decouple activity from expression, and activity may thus add to the analysis of the function of this complex. In cases where miRNA expression is measured at the premature level, activity calculation could provide insight into the usage of the mature strands. For instance, for miR-122 it is clear that 5p is the active strand.

We showed by downsampling that good correlations between expression and activity were obtainable with read counts well below current standards for scRNAseq libraries. We further demonstrated the use of the method on two scRNAseq datasets in mouse and human and with relative high and low read coverage. This demonstrates that the method may be used to infer miRNA activity at the single cell level, and as such should be a useful hypothesis generating tool for studying miRNAs.

Importantly, this type of analysis applied to large datasets could easily be extended beyond miRNAs and 3’UTR sequences. A close analogy lies in transcription factors binding to promoter sequences and RNA binding proteins binding to 3’UTRs. This type of approach may thus facilitate studies of regulatory mechanisms in single cells.

## Methods

### Single cell expression data

For both single cell data sets, we used the following procedure to create inputs for miRNA activity calculations: Cells with less than 1,000 mapped reads across all genes were removed. Counts were divided by the total count in each cell and multiplied by the max total count across all cells. Counts were added a pseudocount of one and logged (base 2).

### 3’UTR sequences

We used 3’UTR sequences as provided by GENCODE v. 19. If genes had multiple transcripts with different 3’UTR lengths, we used the longest form. We removed genes with 3’UTR lengths less than 20 bases and longer than 10,000 bases.

### Fold change value calculations

We used logged expression values as described above, and calculated a fold change value for each gene in each sample or cell by subtracting an expected expression value for that gene. The expected expression value was calculated as the median expression value across all samples or cells in an analysis run, e.g. for TCGA this would be the median expression value across all 9,679 samples.

### miRNA binding model

For all miRNAs, we used the 7-mer sequence complementary to the miRNA seed site as the motif defining miRNA binding sites. We used miRNA mature sequences as defined in miRbase v.20^17^ and used bases from position 2 to 8 from the 5’end as the seed site definition.

### Probability of motifs in sequences

For each 7-mer miRNA binding model, we used Regmex to calculate probabilities for observing the 7-mer motif in sequences similar to the 3’UTR sequence of each gene. Such probabilities were harvested for all 7-mers and 3’UTRs.

### miRNA activity calculation

For each sample or cell, we ordered genes by their fold change values. We next calculated miRNA activity from the probability output of three different motif enrichment tools, Regmex^8^ in brownian bridge as well as modified rank sum mode and Sylamer^9^. We used the ordered 3’UTR sequences as input for Sylamer. Similarly for the Regmex methods, we used the ordered 3’UTR sequences and pre-calculated motif probabilities in the sequences as input, and calculated miRNA activity scores as -log10(p)*sign(statistics).

We provide a tool, miReact, available at https://github.com/muhligs/miReact that implements the method using the modified rank sum mode for use on large expression data sets. See Supplemental note for a tutorial of running miReact on the Tabula Muris data set.

## Data availability

The ‘Tabula Muris’ data set is available at (https://doi.org/10.6084/m9.figshare.5829687.v7).

The human liver data set is available in the Gene Expression Omnibus (GEO) with the accession code GSE124395.

## Code availability

The miReact software is written in R and available under the MIT license at https://github.com/muhligs/miReact.

