## Supplemental Figures for "miRNA activity inferred from single cell mRNA expression"

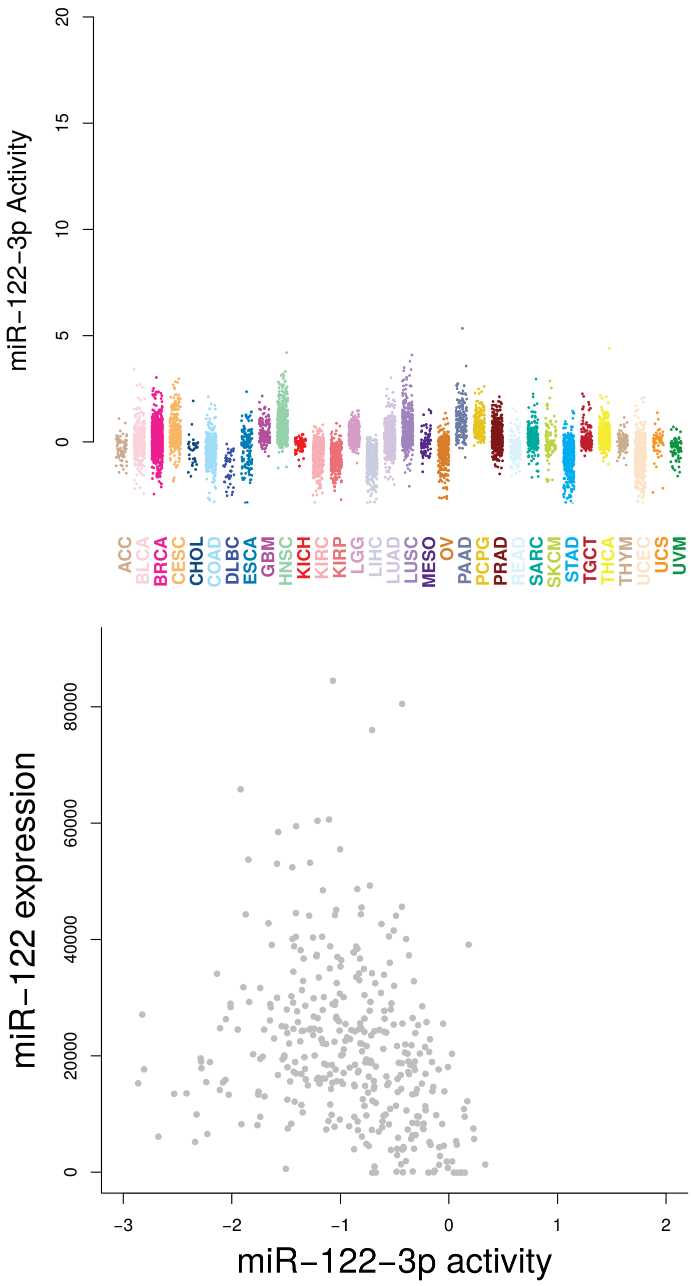


**Supplemental Fig 1: miR-122-3p activity. a,** miR-122-3p activity in 9,679 cancer samples from the TCGA project divided into their primary tissue origin. **b,** Lack of correlation between miR-122-3p activity and miR-122 expression.

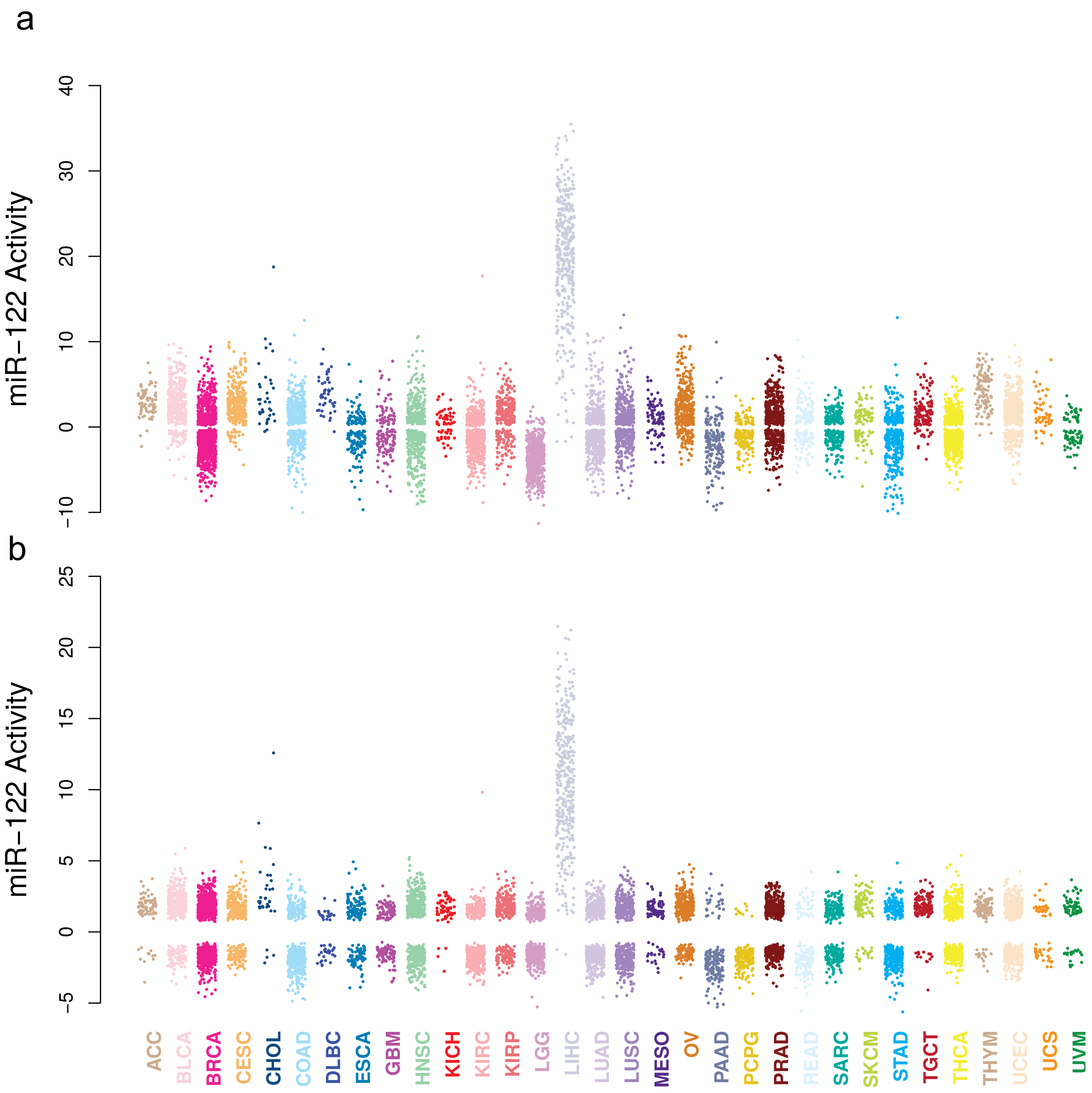


**Supplemental Fig 2: mir-122-5p activity for bb and Sylamer method.** miR-122-5p activity in 9,679 cancer samples from the TCGA project divided into their primary tissue origin. **a,** Activity calculated with the bb method. **b,** Activity calculated with the Sylamer method.

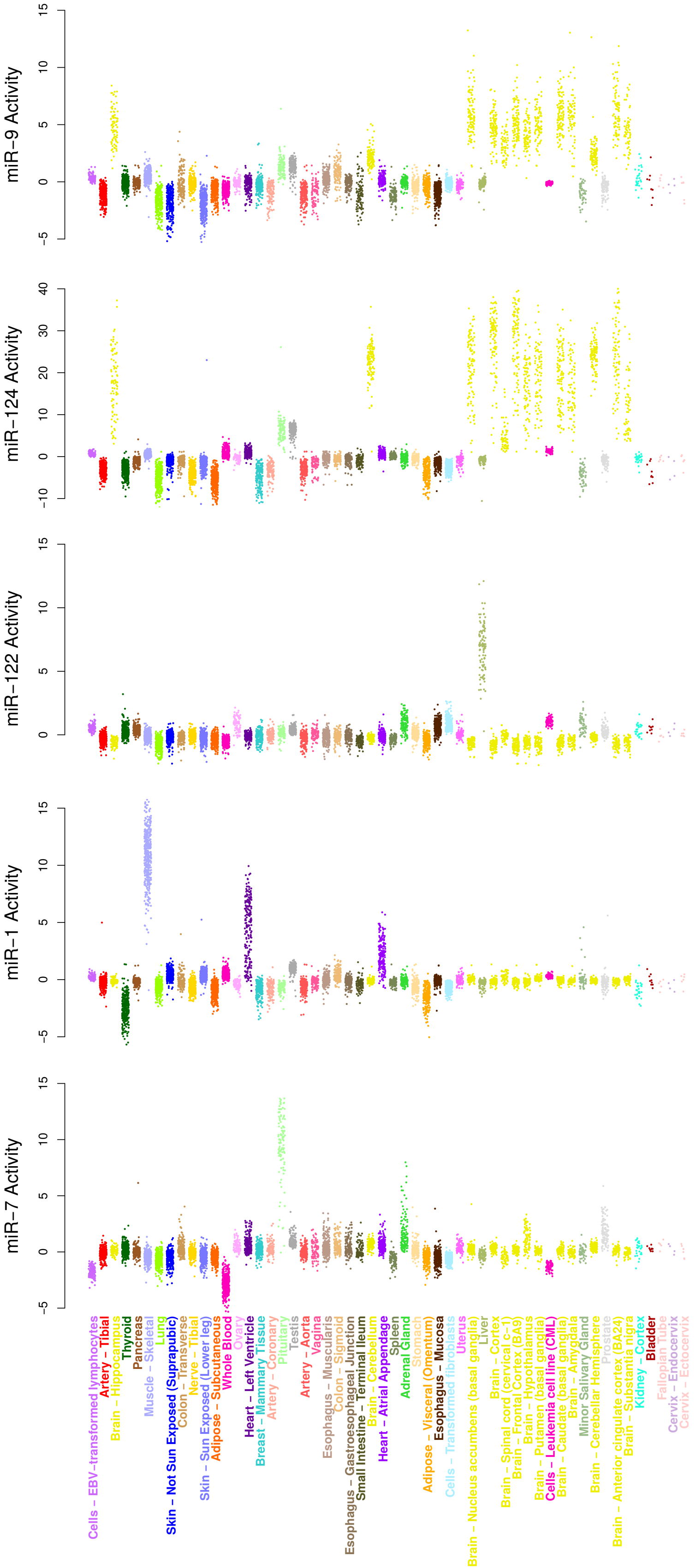


**Supplemental Fig 3: Activity of miRNAs in GTEx tissue data set.** Activity of miRNAs in 7,862 tissue samples from the GTEx project. All runs were made with the mrs method.

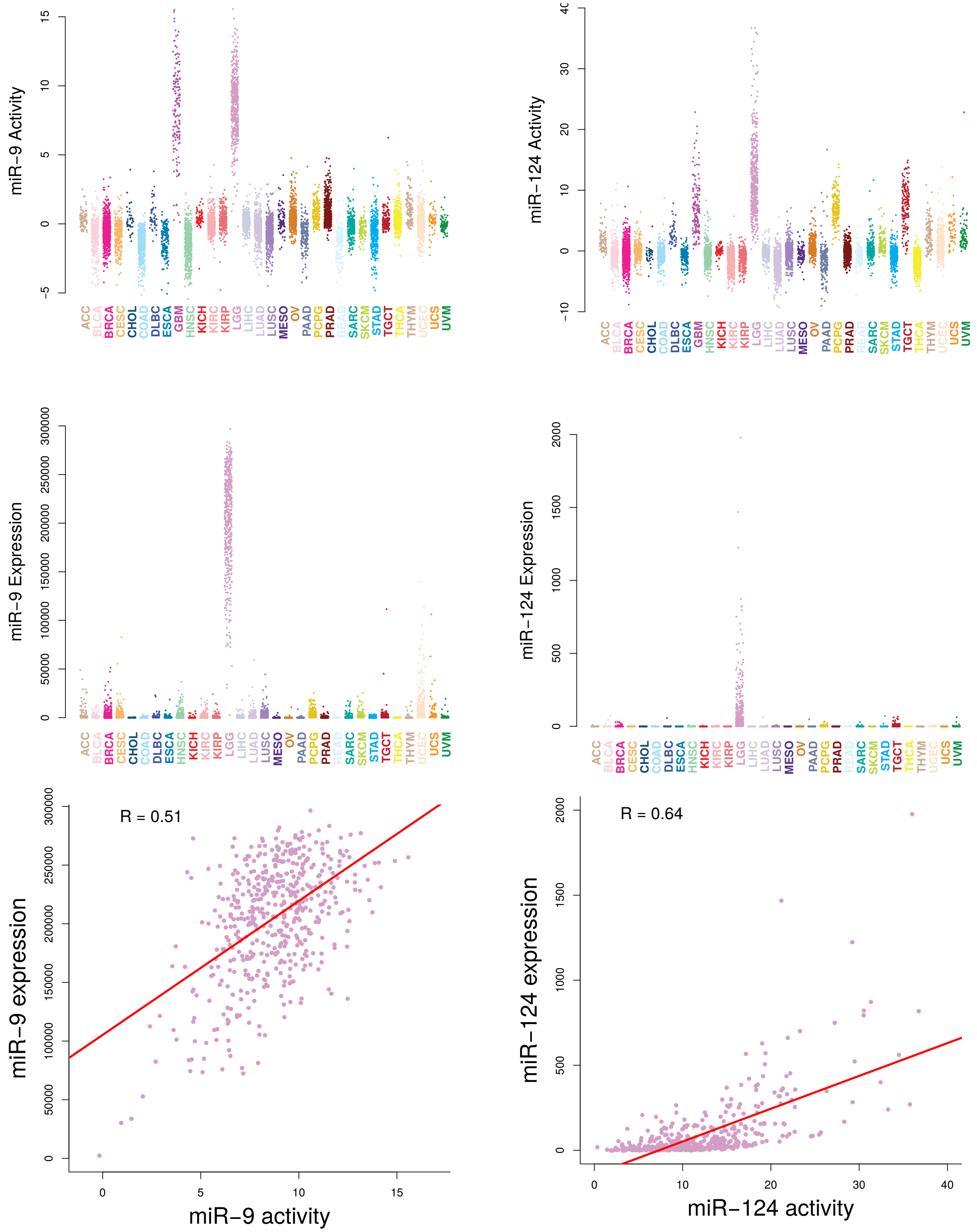


**Supplemental Fig 4: miR-9 and miR-124 expression and activity. a,b,** miR-9 and miR-124 activity in 9,679 cancer samples from the TCGA project divided into their primary tissue origin. **c,d,** miR-9 and miR-124 expression in the same samples excluding GBM for which miRNA expression was not obtainable. **e,f,** Correlation between expression and activity for mir-9 and mir-124 in lower grade glioma samples.

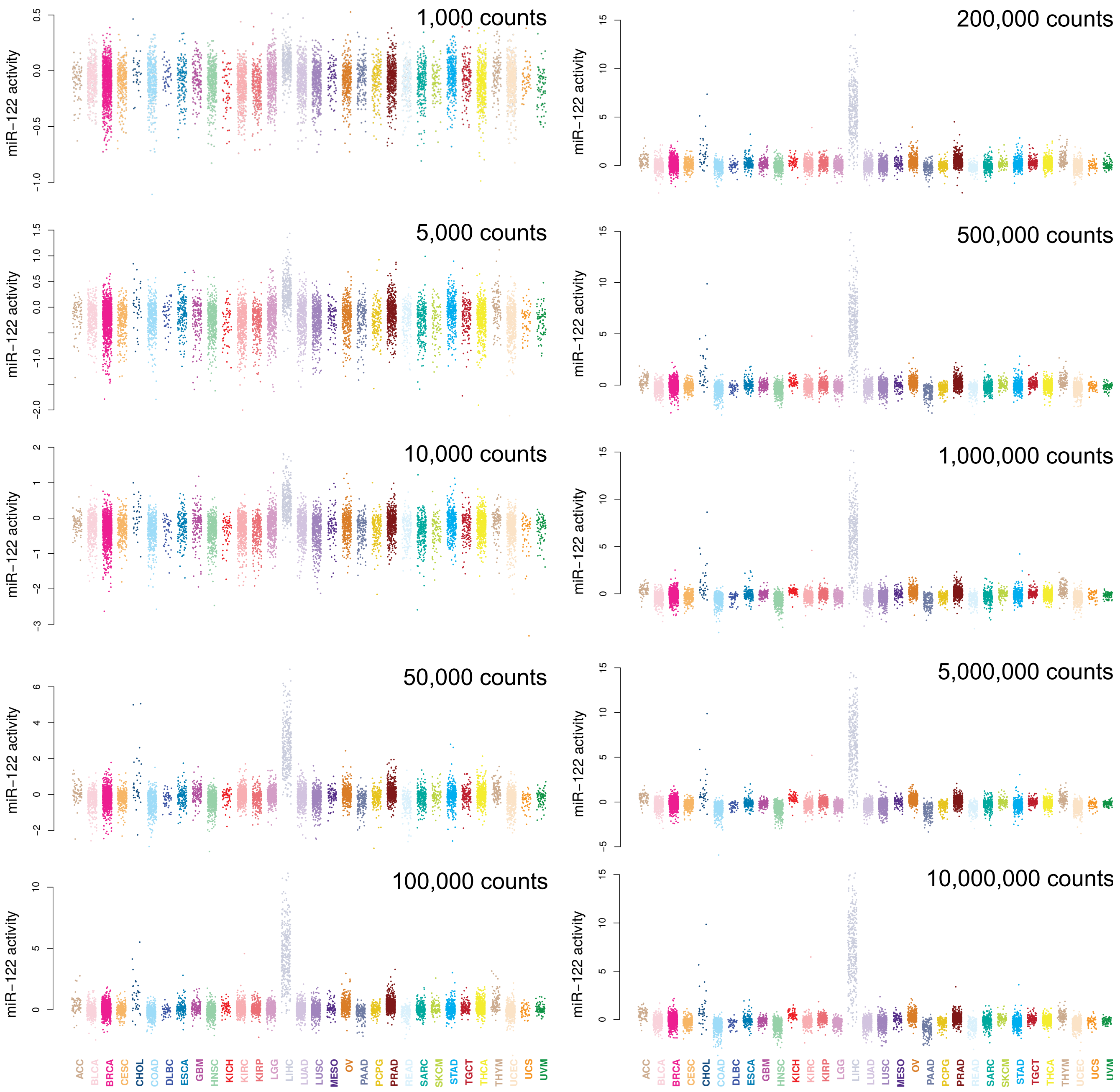


**Supplemental Fig 5: Downsampling activity.** Activity of miR-122 in samples from the TCGA project which were subject to downsampling. Downsampled expression counts for each sample was made by sampling the indicated number of read counts proportional to the samples measured gene expression.

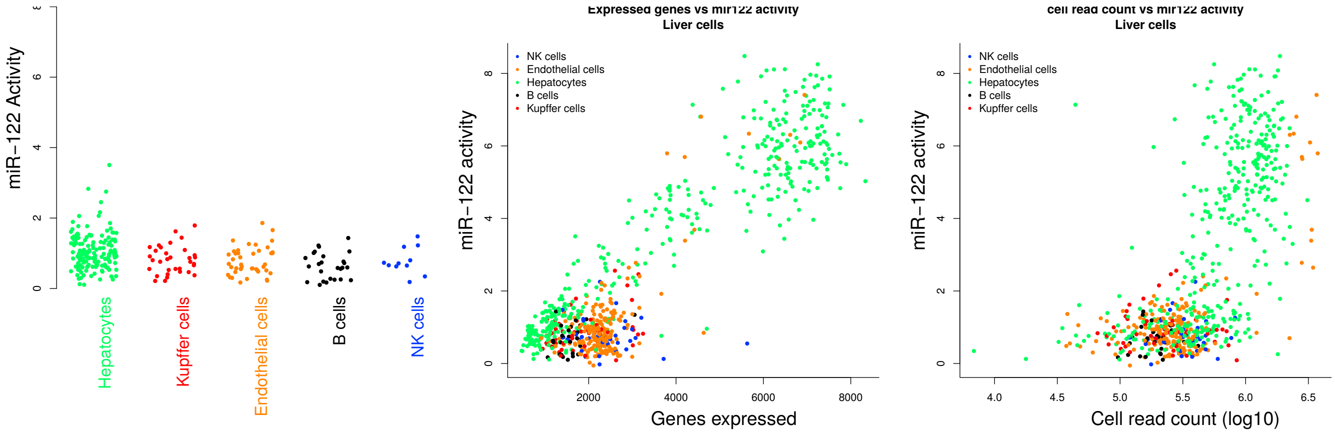


**Supplemental Fig 6: Read count dependence on miR-122 activity in mouse liver cells. a,** Activity of miR-122 in cells with less than 2,000 genes expressed. **b,c,** Effect of miR-122 activity estimates on number of genes expressed and number of read counts for each cell.

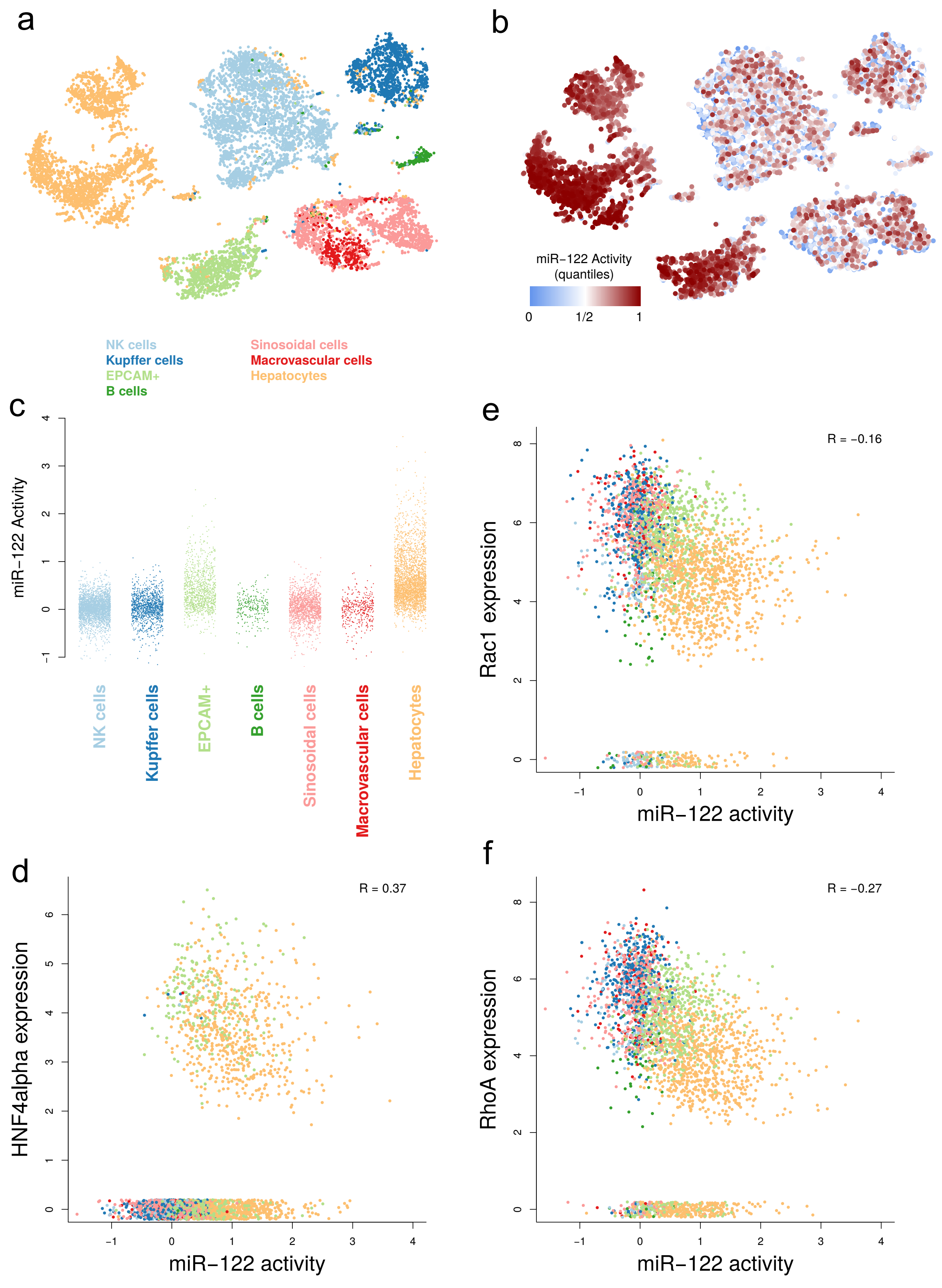


**Supplemental Fig 7: miRNA activity in human liver single cells. a,** t-SNE plot showing clusters of human liver cell types. **b,** miR-122 activity overlaid the t-SNE embeddings as in (**a**). Color scale is based on miR-122 activity quantiles. **c,** miR-122 activity of individual liver cells separated into cell types. **d,e,f,** Correlation between mir-122 activity and expression of HNF4alpha (**d**), Rac1(**e**), and RhoA (**f**). Cells with less than 2000 expressed genes were removed in **d-f**.

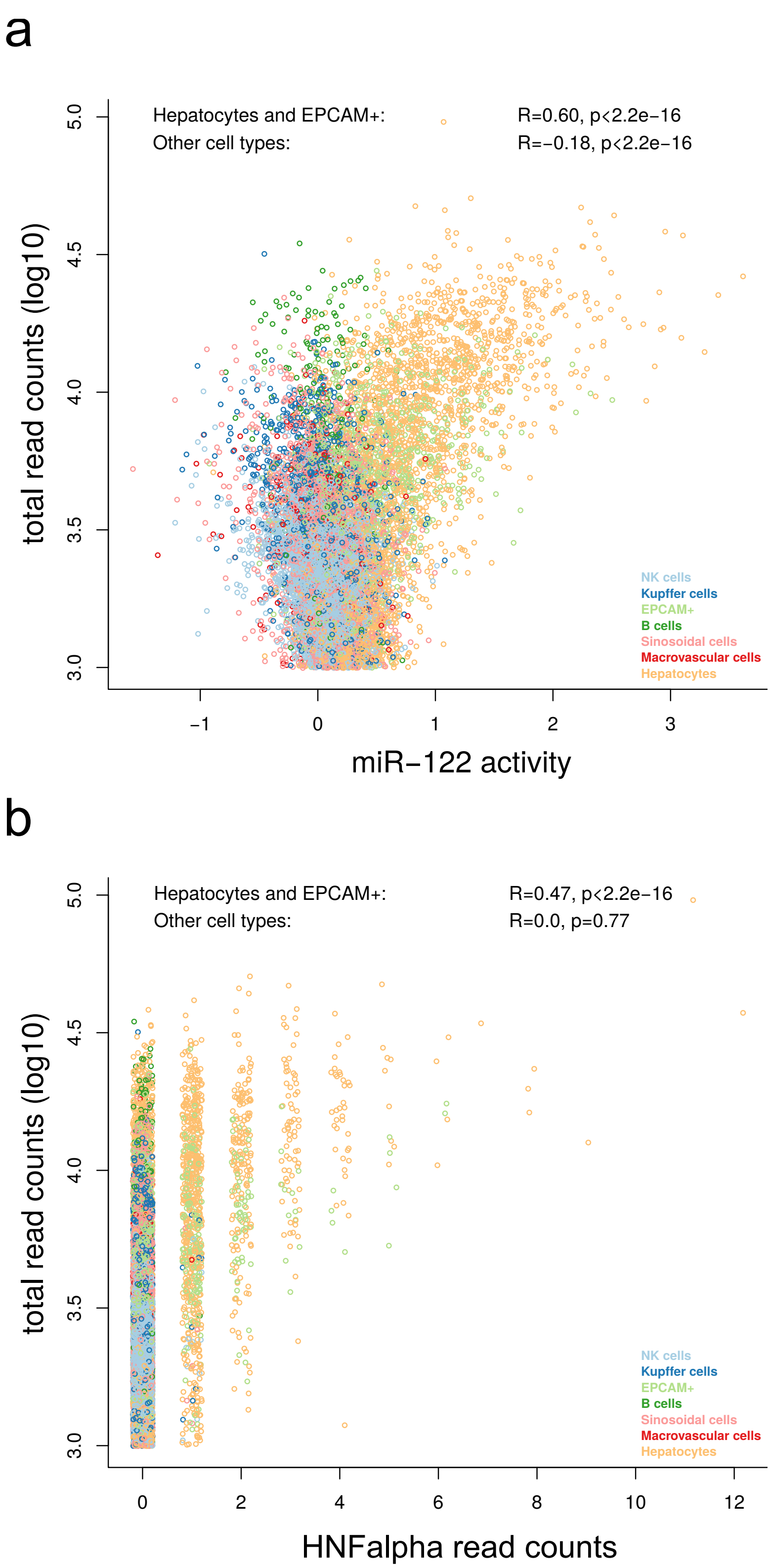


**Supplemental Fig 8: Read count dependence on miR-122 activity and HNF4alpha expression in human liver cells. a,** Effect of miR-122 activity estimates on number of read counts for each cell **b,**Correlation of HNF4alpha read counts with total read counts in hepatocytes and EPCAM+ cells. Failure to capture true expression in low read count cells may explain why some hepatocytes show miR-122 activity without HNF4alpha expression.

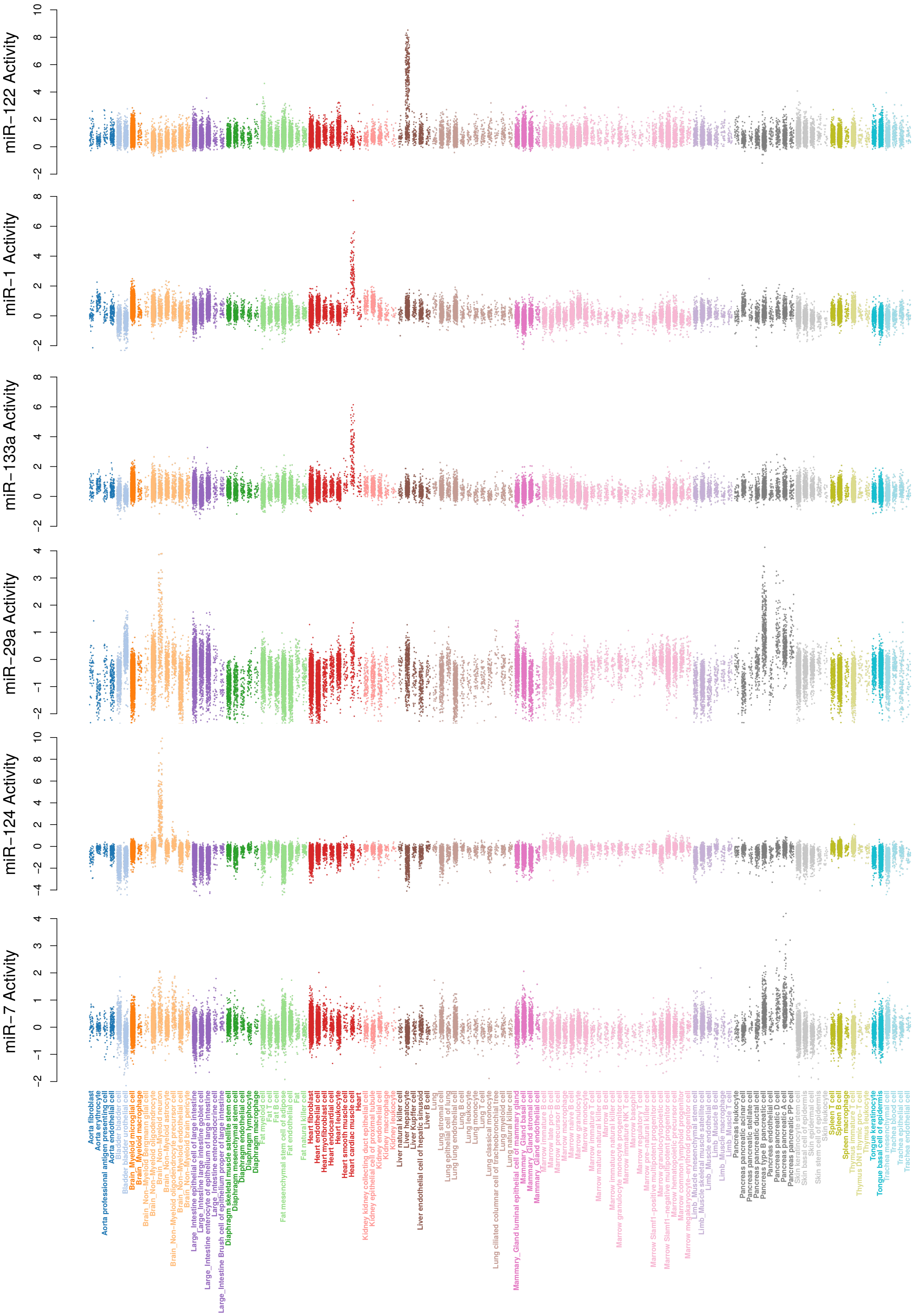


**Supplemental Fig 9: Activity of miRNAs in the Tabula Muris data set.** Activity of miRNAs in 42,192 cells from the mouse single cell data set. Cells were divided into tissues and cell types annotations obtained from the original work.
