## Supplemental Note for "miRNA activity inferred from single cell mRNA expression"

### miReact tutorial

#### Running miReact on Tabula Muris scRNAseq data set

This tutorial will walk through the steps required to produce miRNA activities from the Tabula Muris FACS sorted data set consisting of mRNA expression measures of ~42,000 cells from 18 tissues.

The end result will consist of a matrix of all 16,384 7-mer motifs evaluated for all ~42,000 cells. A subset of the motifs corresponds to miRNA targets by being complimentary to the miRNA seed sites.

The procedure needs to be run on a cumpute cluster with the sbatch queueing system installed.

For convenience, the miReact repository contains a 1000 cell sampled set of the full data set.

##### Obtaining the miReact software

miReact is available at the github repository at:

https://github.com/muhligs/miReact

You can either use the Git program to clone the directory, or download the directory from the web.

In the following, we assume that you will place the directory on a compute cluster (required) in the ~/temp directory, so that the directory ~/temp/miReact contains the directories code, data, motif.models and seqs. If miReact is placed differently, please adjust the code in the tutorial accordingly.

The software depends on the R packages 'Regmex', 'expm' and 'parallel', so these should also be installed.

In R do:

```
###############################################
# install expm
install.packages("expm")
# install devtools (if needed) to install Regmex from github:
install.packages("devtools")
library(devtools)
install_github("muhligs/Regmex")
###############################################
```

##### Calculating motif probabilities in sequences

Prior to making a miReact run, we will need to calculate probabilities for observing motifs in sequences.

This is a somewhat heavy process, and requires multiple cores in an R process. For the mouse sequences, on a laptop/desktop with 16 GB of ram and use of 8 cores, it takes around 10 hours.

If you have more cores available on a cluster node, it is advisable to use it. We have run this on up to 36 cores in less than two hours, but this may require >128 GB of memory.

In addition we need a count matrix of the motifs in the sequences.

In R we do:

```
################################################
# Calculating motif probabilities in sequences
setwd("~/temp/miReact")
# required packages
require(parallel)
require(Regmex)
require(expm)
source("./code/lambdaProbdist.R") # functions required
seqlist <- readRDS("./seqs/mm.utr3.seqlist.rds") # this is the sequences for mouse 3'UTRs.
load("./motif.models/patterns.7mer.Rdata") # these are motif models

# produce and save the probability matrix
# adjust the number of cores accordingly (mc.cores =)
reslist <- mclapply(patterns,
  function(x)unlist(lapply(seqlist,function(y)pd.mrs2(x,y))),mc.cores = 8)
names(reslist) <- Regmex::all.mers(7)[1:length(reslist)]
pval.mat <- do.call(rbind,reslist)
pval.mat[1:5,1:5] # motifs x sequences
dim(pval.mat) # 16,384 motifs x 52,419 3'UTR sequences
saveRDS(pval.mat, file="./motif.models/mm.seqXmot.utr3_mrs_7mer.rds")

# produce and save the count matrix
# adjust the number of cores accordingly (mc.cores =)
counts <- do.call(rbind,mclapply(Regmex::all.mers(7),
  function(x)Regmex:::n.obs.mot(x,seqlist,overlap=FALSE),mc.cores=8))
rownames(counts) <- Regmex::all.mers(7)
dim(counts) # 16,384 motifs x 52,419 3'UTR sequences
seqs <- readRDS("./seqs/mm.utr3.seqs.rds")
colnames(counts) <- seqs$tid
saveRDS(counts, file="./motif.models/mm.seqXmot.counts.utr3_mrs_7mer.rds")
################################################
```

These files are also available at the Synapse repository at:

https://www.synapse.org/#!Synapse:syn22227236

Download these files ('mm.' for mouse runs and 'hs.' for human runs) and place them in the miReact/motif.models directory.

##### Obtaining and preparing the Tabula Muris data

The Tabula Muris data set can be obtained from

https://figshare.com/articles/Single-cell_RNA-seq_data_from_Smart-seq2_sequencing_of_FACS_sorted_cells/5715040.

Download the FACS.zip file and unpack into ./data/tm

This should produce 18 .csv files corresponding to 18 different tissues.

In R, prepare the expression data set with the following commands:

```
################################################
# make raw count expression matrix
setwd("~/temp/miReact/data")
lf <- list.files("tm",full.names = T, pattern = "counts.csv")
```

```r
d <- read.table(as.is=TRUE, sep=",", quote = "", header=TRUE, lf[1],row.names = "X..")
dim(d) # 23,433 x 1,638

for(i in lf[2:length(lf)]){
  d <- cbind(d,
  read.table(as.is=TRUE, sep=",", quote = "", header=TRUE, i,row.names = "X.."))
  print(paste(i,dim(d)[2]))
}
dim(d) # 23,433 x 53,760
# save raw expression matrix
saveRDS(d, file="tm_exp_raw.rds")
###############################################
# make an exp file for miReact...
seqs <- readRDS("~/temp/miReact/seqs/mm.utr3.seqs.rds")
idx <- match(gsub("\"","",rownames(d)),seqs$gsym)
# remove genes without 3'UTRs from expression matrix
d <- d[!is.na(idx),]
# rename genes to ensemble geneIDs
rownames(d) <- seqs$gid[idx[!is.na(idx)]]
dim(d) # 18,005 x 53,760 # 18,005 genes used in the analysis
# define library counts
libs <- apply(d,2,sum)
hist(log10(libs),100) # look at distribution of library counts
saveRDS(libs, file="tm_libs_count.rds")
min(libs) # 14
max(libs) # 12,450,458
median(libs) # 426,493.5
# remove cells with less than 1000 reads or more than 5,000,000 reads
sum(libs<1000) # 3795
sum(libs>5000000) # 27
d <- as.matrix(d)
d <- d[,libs>1000&libs<5000000]
median(libs[libs>1000&libs<5000000]) # 464,902
dim(d) # 18,005 genes x 49,935 cells
# normalize colwize expression to 1 and multiply by median
d <- sweep(d, 2, colSums(d), FUN = '/')*median(colSums(d))
d <- log2(d+1) # log transform
saveRDS(d, file="tm_mireact_exp.rds")
###############################################
```

##### Running miReact on Tabula Muris data

The miReact run should take place on a compute cluster with the sbatch queuing system installed.

The program will start a process for every 20 samples/cells, and thus this data set will spawn around 2,000 processes.

We have prepared a smaller data set with 1000 cells available on Synapse at:

https://www.synapse.org/#!Synapse:syn22227230

That can be used to test the procedure before performing a full run.

Place this file in the data directory, i.e. ~/temp/miReact/data/mm.exp1000downsample.rds

The mireact function creates a working directory under ~/temp/miReact with format:

o[currentDateAndTime]

Inside this directory, there is an

Rscript-[processID].out

file where the process of the run can be followed.

If the miReact software was placed in a different directory than ~/temp this should be stated in the install.dir parameter (install.dir = "path/to/directory")

in R do:

```
################################################
# Run mireact
source("~/temp/miReact/code/mireact.R")
efile <- "~/temp/miReact/data/tm_mireact_exp.rds"
# If running on the smaller test set, do:
efile <- "~/temp/miReact/data/mm.exp1000downsample.rds"
# This following command starts miReact.

# A mail message can be optionally created
# if the 'mail' program is installed on your system.
# This will send a mail at the start and end of the run.
# Please unset this mail parameter (default mail = NULL) if not available or desired.

# If the miReact software was placed in a different directory than ~/temp
# this should be stated in the install.dir parameter.
# Adjust the name of the outputfile (out.file) accordingly.

mireact(exp=efile, species="mm", motifs=7, seq.type="utr3",
out.file="~/temp/miReact/data/mirnaActivity_tm_1000.rds",out.meonly=T,
mail="", install.dir = "~/temp/miReact")
################################################
```

Now make some plots of the caluclated miRNA activities. Here shown for the reduced 1000 cell data set. Adjust filenames if needed.

```
################################################
# load motif activity matrix
ma <- readRDS("~/temp/miReact/data/mirnaActivity_tm_1000.rds")
# load cell annotations for sample set
a <- readRDS("~/temp/miReact/data/mm.annotations1000downsample.rds")

## FULL SET ONLY ##########
# load full set annotations
a <- readRDS("~/temp/miReact/data/tm.annotation.rds")
# make ma follow annotation order (and contain cells with annotations only)
colnames(ma) <- sub("[.]$","",sub("X.","",colnames(ma)))
ma <- ma[,a$cell]
## END FULL SET ONLY ######

# source functions
source("~/temp/miReact/code/plot.tools.R")

# plot t-SNE of cells colored by tissue
plot(a$tsne1, a$tsne2, col=a$color, main="", pch=20, cex=1, xlab="", ylab="", axes=F)
text(centroid(a[,c("tsne1","tsne2")], a$cluster2),
```

```
  labels=a$clusternaming[match(unique(a$cluster2), a$cluster2)],
  cex=1, col=1, lheight=0.1, font=2)
  legend(par("usr")[1]+10, par("usr")[3]+4, unique(a$tissue2), col=unique(a$color),
         bty="n", pch="", ncol = 4, cex=0.7, pt.cex = 0,
         text.col = unique(a$color), xpd=T, text.font = 2)
# now plot the activity overlay for miR-122
plot(a$tsne1[order(ma["ACACTCC",])], a$tsne2[order(ma["ACACTCC",])],
  col=colorme(ma["ACACTCC",], ramp = c("cadetblue1", "white", "darkred"),
  numcol=99, quantcol=F)[order(ma["ACACTCC",])], pch=20, cex=1.5,
  xlab="", ylab="", axes=F, main="miR-122-5p activity")
setKey(colorRampPalette(c("cadetblue1", "white", "darkred"))(99), title="",
  pos=c(-0.05,0.06), label=c("low", "", "high"), height=0.05, keylength = 0.25)

# plot the activities as dots
par(mar=c(9.1,5.1,4.1,2.1))
bardot(ma["ACACTCC",], a$tissue2,col = a$color, main="", ylab="miR-122 Activity",
  ylims=c(-1.1,9), labelpos = -0.03, cex=0.1, labelcex=1,
  labeladj=0.45, ylabcex=1.5, labfont=2)
##############################################
```
